# Electroencephalography signals in a female Fragile X Syndrome mouse model

**DOI:** 10.1101/2024.04.04.588163

**Authors:** Asim Ahmed, Veronica Rasheva, MoonYoung Bae, Kartikeya Murari, Ning Cheng

## Abstract

**Background:** Fragile X syndrome (FXS) is the leading monogenic cause of Autism. No broadly effective support option currently exists for FXS, and drug development has suffered many failures in clinical trials based on promising preclinical findings. Thus, effective translational biomarkers of treatment outcomes are needed. Recently, electroencephalography (EEG) has been proposed as a translational biomarker in FXS. Recent years have seen an exciting emergence of novel EEG signal analyses from FXS patients. However, there is a notable gap in corresponding analyses conducted on animal models of the disorder. Being X-linked, FXS is more prevalent in males than females, and there exist significant phenotype differences between males and females with FXS. Recent studies involving male FXS participants and rodent models have identified an increase in absolute gamma EEG power, while alpha power is found to be either decreased or unchanged. However, there is not enough research on female FXS patients or models. In addition, studying EEG activity in both young and adult FXS patients or rodent models is crucial for better understanding of the disorder’s effects on brain development. Therefore, using the well established *fmr1* knockout (KO) mouse model of FXS, we aim to compare EEG signal between female wild-type (WT) and female model mice at both juvenile and adult ages.

**Methods:** Frontal-parietal differential EEG was recorded using a stand-alone Open-Source Electrophysiology Recording system for Rodents (OSERR). EEG activity was recorded in three different conditions: a) in the subject’s home cage, and in the arenas for b) light -dark test and c) open field test. Absolute and relative EEG power as well as peak alpha frequency, theta-beta ratio, phase-amplitude and amplitude-amplitude coupling, and EEG signal complexity were computed for each condition.

**Results:** In our study, we found absolute alpha, beta, gamma and total EEG power is increased in the female model compared to WT controls at the juvenile and adult ages. Alongside, relative theta power is decreased in the model. Additionally, phase-amplitude and amplitude-amplitude coupling is altered in the model. Furthermore, peak alpha frequency is increase, and theta-beta ratio is decreased in the model. Lastly, no change in EEG signal complexity is found.

**Discussion and Conclusion:** Consistent with most findings from FXS patients and rodent models, our results demonstrated an increase in gamma power in *fmr1*KO female mice, reinforcing gamma power as a robust and reliable EEG phenotype across FXS models. Additionally, theta-gamma cross frequency amplitude coupling is inversely coupled in female FXS model, which is similar to what has been reported in FXS patients. Overall, our findings reveal that not all EEG biomarkers observed in FXS patients are replicated in the female FXS model. For example, peak alpha frequency, theta-beta ratio, and brain signal complexity showed discrepancies between the mouse models and FXS patients. Additionally, when compared to previously reported EEG changes in male FXS mouse models, our results highlight the presence of a potential sex-based difference in EEG phenotypes at both juvenile and adult stages of *fmr1* KO mouse models. Together, our study indicates that certain EEG parameters may be more translatable between rodent models and FXS patients than others and underscore the importance of considering sex and developmental stage as a critical factor when using EEG as a biomarker in FXS research.

## Introduction and Background

Autism is one of the most common neurodevelopmental disorders worldwide (1, 2). The leading monogenic cause of autism and intellectual disability is fragile X syndrome (FXS). The fragile X messenger ribonucleoprotein (FMRP), encoded by the *fmr1* gene, is lost due to mutations in FXS (3, 4). FXS is estimated to occur 1 in 4,000 males and 1 in 7,000 females (5). There is no broadly effective support option for autism or FXS, and numerous clinical trials of drugs based on promising preclinical results have failed (6-9). Therefore, it is necessary to develop accurate translational biomarkers, allowing for more fruitful clinical tests and translational success.

Recently, electroencephalography (EEG) has been identified as a potential translational biomarker of neurodevelopmental disorders, as it can record neural activities similarly in humans and animals (6-10). Notably, spontaneous EEG data from male FXS patients and rodent models indicate increased power in the gamma band (11). Therefore, increased gamma power is suggested to be a transferable hallmark across FXS patients and animal models.

EEG is a method of *in-vivo* electrophysiological recording, and it measures electrical activity from the cerebral cortex. EEG signals are generated by the collective electrical activity of neurons in the brain, especially due to the synaptic activity of cortical pyramidal neurons. The rhythmic and synchronized firing of large groups of neurons leads to detectable electrical potential on the scalp, captured by electrodes. This process involves the spatial summation and temporal coherence of neural oscillations (12). EEG signal comprises low-frequency bands including delta (1-4Hz), theta (4-8Hz), and alpha (8-13Hz), and high-frequency bands, beta (13-30Hz), and Gamma (30-100 Hz) (13). It is believed that the slow waves of the low-frequency bands are essential for long-distance or interregional communication, whereas the fast waves of the high-frequency bands are important for short-distance or intraregional communication (13).

The *fmr1* knockout (KO) mouse model has been employed for over two decades as a fundamental preclinical model for studying FXS (14). The absence of the FMRP protein in these mice leads to changes in neuronal structure, such as altered spine shape and density (15, 16), as well as behavioral phenotypes including heightened locomotor activity and increased sensitivity to sensory inputs (17, 18).

Being an X-linked disorder, FXS is more prevalent in males compared to females (4, 5). Females also have FXS, but homozygous mutation is rare. Heterozygous mutation in females manifests milder symptoms, possibly due to skewed X chromosome inactivation (1). In one study of adult FXS patients, male and female participants show an increase in baseline gamma and theta power and decrease in alpha power in EEG, and no sex difference was reported (11). Another study in adult FXS patients reported a sex difference, where FXS male patients showed an increase gamma power, while FXS female patients showed decrease gamma power. However, both sexes showed similar EEG phenotypes of increased relative theta, and decreased beta power in FXS groups (19).

In rodent models, male mice also show an increase in spontaneous gamma power in both juvenile and adult age (5, 20). In rat models, adult *fmr1* KO males showed reduced alpha power and increased gamma power (21).

In general, there is very limited research with female models of FXS. One study with adult female *fmr1* KO mice on B6 background reported no difference in gamma EEG power compared to wildtype controls. However, findings in other frequency bands are not reported (22). Another study in adult female *fmr1* KO rats found decreased relative theta power (23). Therefore, understanding of the EEG phenotypes in rodent models of FXS has been incomplete, especially regarding females and their developmental changes.

Recently, new alterations in resting-state EEG spectral domains have been identified in FXS and other neurodevelopmental conditions, which may serve as specific EEG biomarkers reflecting brain maturation, network dynamics and hyperexcitability (24). Specifically, peak alpha frequency, theta-beta ratio, phase amplitude and amplitude-amplitude coupling, and signal complexity have been proposed as sensitive indicators of atypical brain maturation, cortical hyperexcitability and the presence of neurodevelopmental disorders (24, 25). However, they have not been well studied in animal models, and it remained unclear whether they represent consistent phenotypes across patients and models.

In the current study, our objective was to investigate EEG phenotypes in the female *fmr1* KO mouse model at both juvenile (P40) and adult (P70) ages. To this end, we conducted EEG recordings from mice at P40 and again at P70 and compared KO and WT genotypes. Our results indicated that absolute alpha, beta, and gamma EEG power was increased in the KO mice compared to WT controls at both P40 and P70. Alongside, relative theta power was decreased in the KO mice. Additionally, markers of network dynamics and hyperexcitability, including phase-amplitude and amplitude-amplitude coupling was altered in KO mice. Furthermore, EEG parameters reflecting brain maturation and cognition including peak alpha frequency and theta-beta power ratio were also affected in KO mice, while signal complexity remained unchanged.

## Materials and Methods

### Animals

Breeder wildtype (FVB.129P2-Pde6b+ Tyrc-ch/AntJ, Jax stock No: 002848) and *fmr1* knockout (*fmr1 KO*-fmr-tm1Cgr, FVB.129 Jax stock No: 004624) mice were obtained from the Jackson Laboratory (ME, USA) and maintained at the mouse facility of the Cumming School of Medicine, University of Calgary. EEG testing was performed on female KO and WT mice, bred in our animal facility at the University of Calgary. All *fmr1* KO mice of the current study were generated by mating *fmr1* KO (*fmr1* -/-) homozygous females with *fmr1* KO (fmr1 -/y) males. Mice were group housed (up to five per cage) with their same-sex littermates. The cages were in a room with a controlled 12-hr light-dark cycle (lights on at 7:00am) with access to food and water ad libitum. Mouse pups were weaned around 20 days of age. All mice were fed standard mouse chow. EEG testing was performed between 09:00h and 19:00h.

### Ethics

All procedures in this study were performed in accordance with the recommendations in the Canadian Council for Animal Care. The protocol of this study was approved by the Health Sciences Animal Care Committee of the University of Calgary.

### EEG surgery, recording and Data analysis

EEG surgery was carried out in mice utilizing established techniques (10, 26, 27). Mice were given 5% isoflurane to initiate anesthesia, which was subsequently sustained with 1%–2% isoflurane. Mice were placed on top of a heated heating pad on the stereotaxic equipment. The eyes were covered with eye gels to keep them from drying out. After shaving, the scalp was cleaned with 70% ethanol and iodine. Once it was determined that there was no pinch reflex in the toes, the top layer of skin was removed, and the muscles and connective tissue were carefully pushed aside to reveal the skull. To ensure no connective tissue is left, the skull was further cleaned by hydrogen peroxide. To obtain differential frontal-parietal recordings, three small holes were drilled into the skull. The following coordinates were used to fix the electrodes into the holes: ground (AP: +2.5 mm, ML: +1.3 mm), the left parietal cortex (AP: −2.2 mm, ML: −2.5 mm), and the left frontal cortex (AP: +2.5 mm, ML: −1.3 mm, relative to bregma). After connecting the electrodes to a miniature connector, dental cement was used to affix the assembly to the skull. Following surgery, mice were given analgesic medication and kept on the heating pad until they showed signs of ambulation. EEG recording was conducted 5-7 days following the surgery (**Figure 1A**). After being briefly sedated with isoflurane inhalation, the subject mouse was connected to a stand-alone EEG device (10) and then returned to its home cage for EEG recording. A 30-minute recording session in the home cage was followed by a 30-minute recording session in the light-dark chamber. Next day, the same recording was conducted except that the home cage recording was followed by recording in the open field chamber (**Figure 1B**). EEG recording was done at juvenile (P40) and adult (P70) ages.

**Figure 1:**
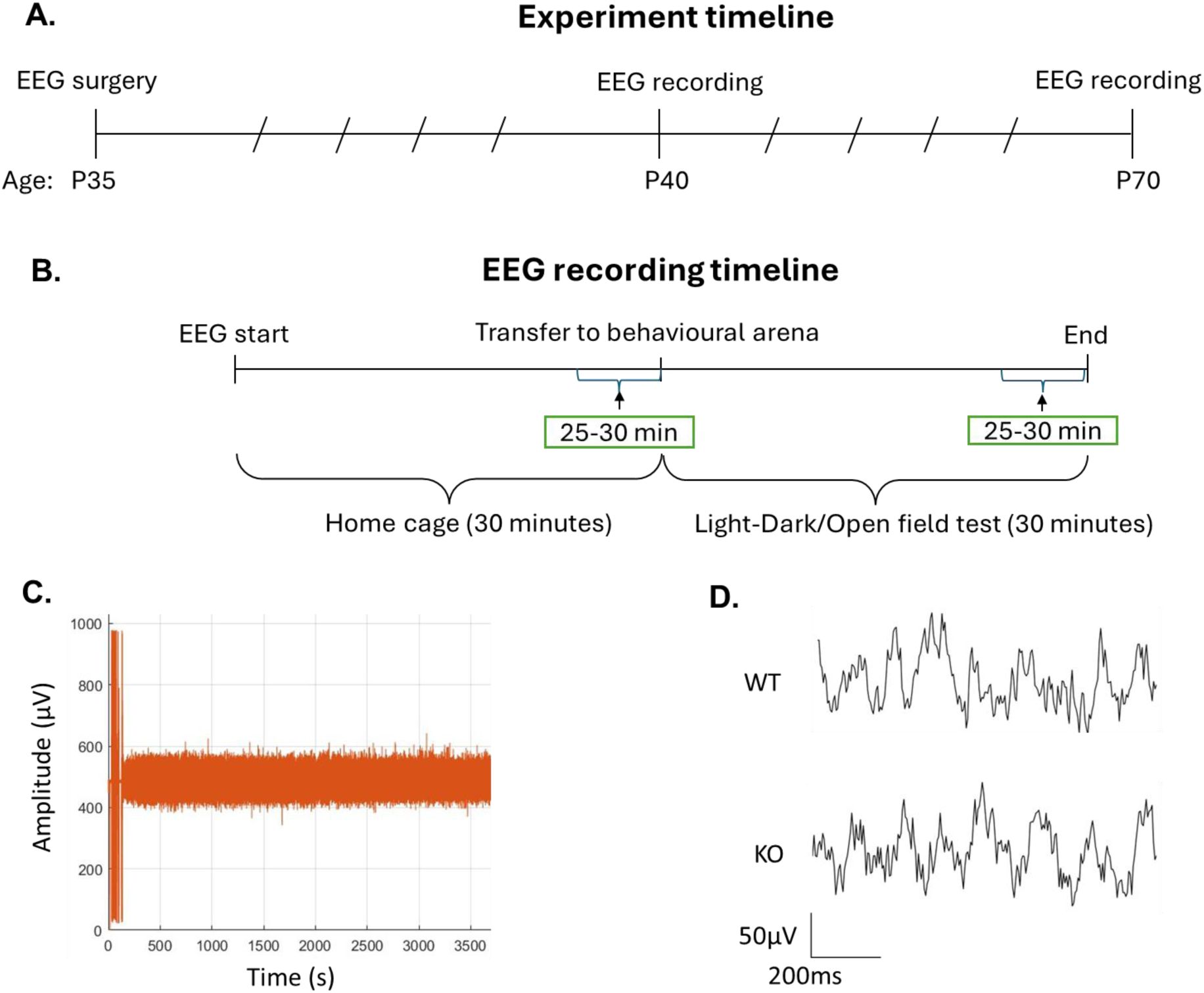
Experimental timeline and EEG recording timeline. (A) Experimental timeline, with EEG surgery at P35 and EEG recording at P40 and P70. (B) EEG recording timeline; first 30 minutes of EEG recording were in the home cage and the next 30 minutes were in the behavioral arena. For analysis, we used around the last 5 minutes (25-30 minutes) of EEG recordings from the home cage and the behavioral arena. (C) EEG raw signal from 1 hour of recording. The noise in the first 120 seconds (about 2 minutes) shows the time when the mouse was anesthetized, and the EEG recording device was connected to the mouse head cap. (D) Example of EEG signal.

Light-dark arena: The apparatus consists of a dark and a brightly lit compartment. The subject was free to move between the two compartments during the 10 minutes test (28).

Open field arena: A subject was placed in an empty square shaped chamber and monitored for 10 minutes under room light (3).

Standard procedures were followed for the analysis of EEG data (26, 27). The recording quality was assessed (**Figure 1C**), and a 5-minute EEG segment around the end of each condition (home cage, Light-Dark, and Open Field) was chosen for analysis. Additionally, we ensured that mice remained active during these EEG segments, by analyzing video recordings. Using MATLAB, Welch’s approach (29) was used to compute the power spectral density (PSD) of the EEG signal. We divided the EEG data into 2-second epochs. A Hamming window was used to compute periodograms for each epoch and then averaged to get the final PSD estimate at a 0.5 Hz spectral resolution. To determine the power in each of the following frequency bands: delta (1–4 Hz), theta (4–8 Hz), alpha (8–13 Hz), beta (13–30 Hz), and gamma (30–100 Hz), the PSD was integrated using the trapezoidal technique. Two sub-divisions were made for the gamma band: “low gamma (30–60 Hz)” and “high gamma (60–100 Hz).” Furthermore, a Fast-Fourier Transform was used to compute a continuous power spectrum, which was displayed on a decibel scale (10*log10). Relative power was calculated as the ratio of the power of a particular band to the total power of all bands summed together.

### Phase amplitude coupling

Phase-amplitude cross-frequency coupling (PAC) was measured by further analysis of the EEG data. The PAC between gamma band (30–100 Hz) and lower frequency oscillations (4– 12 Hz) was measured using a time-frequency based approach (30). This quantified the modulation of the amplitude of gamma oscillation with the phase of the slow rhythm. Using the RID-Rihaczek distribution, the high frequency oscillation’s envelope and the low frequency oscillation’s phase were obtained. The extracted amplitude and phase time series were then used to compute the mean vector length (MVL), which was used to quantify PAC. MVL measures the circular variances by measuring the complex composite signal’s mean magnitude. It is a modulation index used to measure the modulation between the phase and amplitude time series (31). Two minutes of spontaneous EEG activity were examined. Using a step of 1 Hz for low frequencies (referred to as “phase frequency”) from intervals of 4–12 Hz and a step of 2 Hz for high frequencies (referred to as “amplitude frequency”) from intervals of 30–100 Hz, MVL was computed.

### Amplitude – Amplitude coupling

Amplitude – amplitude coupling analysis was conducted to examine the potential dependence between low-frequency power and high-frequency activity. It is used to investigate the potential associations of alpha and theta power with gamma power (11).

We calculated cross-frequency amplitude coupling over a course of time (11). Time series from our EEG recording were segmented into 2-s epochs. Absolute low and high alpha, theta, and low and high gamma power were calculated for each epoch (32). Spearman’s correlation coefficient for each low frequency (theta, low alpha and high alpha) and high frequency (low gamma and high gamma) was calculated using the time series of mean absolute EEG power across 2-s epochs. Fisher’s Z-transform was used to normalize group-wise comparisons (32)..

### Multi-Scale Entropy

Multi-scale entropy (MSE) has been used to measure signal complexity in participants’ EEG while at rest (33). To calculate MSE from our mouse EEG recordings, we followed the established method (33). MSE calculations were performed using the algorithm outlined by (34), which creates multiple time scales by down-sampling the original EEG signal through a process known as coarse-graining (33). The original time scale is segmented into non-overlapping windows, which are subsequently averaged. As the scale (window numbers) increases, the time series become shorter (33). In this study, the coarse-graining process was applied to 2 s epochs of EEG data for each mouse. Later, sample entropy (SampEn), that quantifies the signal variability for each time series by assessing the predictability of amplitude patterns within it, was quantified (35). The pattern length was set to m=2, meaning the algorithm identified matching sequences of two consecutive points in the signal. The tolerance level was set to r=0.5, indicating that amplitude points within 50% of the standard deviation of the time series were included in the analysis. The algorithm then counted the number of matching sequences for m+1 data points, and SampEn was calculated as the negative natural logarithm of the ratio between the total matches for m+1 and m. Finally, MSE values were averaged across all epochs within the 5 min recording for each mouse, resulting in a final MSE score for each time scale from 1 to 40.

### Statistical tests and analysis

Statistical analyses were performed using GraphPad PRISM, version 10 (GraphPad Software, Boston, MA, USA). The sample size for this study was determined based on prior reports and preliminary data to ensure adequate statistical power. Data from animals with noisy signals were excluded from the analysis. Noisy signals were identified by their abnormally large and often saturating amplitudes. These signals were likely caused by connection issues between the electrodes and the connector, which occurred after surgery. These artifacts were considered non-representative of true neural activity and were removed to maintain data quality and reliability. Values were mean± SEM. Levels of significance were: *: p < 0.05, **: p < 0.0l, ***: p < 0.001. For PSD analysis, log power spectrum was plotted for both WT and KO mice at both P40 and P70 ages, for each frequency in 0.5Hz increments from 1 to 100. Subsequently, individual data was averaged and a two-way ANOVA with repeated measures, followed by post hoc multiple pairwise comparisons was used to compare EEG signal between WT and KO mice at both P40 and P70 for absolute and relative power. Before applying a two-way ANOVA repeated measure, assumptions of normality was also tested. Normality was assessed using the Shapiro-Wilk test (p > 0.05). This assumption was satisfied, justifying the use of parametric testing. Three-way ANOVA repeated measure was performed for measuring the timepoint variability in absolute EEG power across four time points in the recording from home cage followed by light-dark and open field arenas. Peak alpha frequency was calculated as the frequency with maximum amplitude between 8 and 13 Hz for each mouse. To calculate it, EEG data was reanalyzed at obtains PSD values with spectral resolution of 0.1 Hz. Following that, peak frequency was identified for each recording. Later, two-way ANOVA with repeated measures was carried out to compare it between WT and KO at both ages. Theta-beta ratio was calculated as the power in the theta frequency band divided by the power in the beta frequency band for each mouse. Theta-beta ratio was compared between KO and WT mice at both ages using two-way ANOVA with repeated measures. For multi-scale entropy (MSE), the complexity index was calculated as area-under-the-curve for scales 1–40 to obtain a general indication of signal complexity. Two-way ANOVA with repeated measures was used to compare WT and KO at both ages. In a follow-up analysis, scales 1–20 and 21–40 were integrated by averaging the MSE scores 1 - 20 and 21 – 40, to obtain a more fine-grained picture of differences in signal complexity between the two genotype groups that were assessed using a subsequent two-way ANOVA with repeated measures, followed by post hoc multiple pairwise comparisons. For phase-amplitude coupling, MVL values, which quantifies phase-amplitude coupling, were compared using a two-way ANOVA with repeated measures followed by post hoc multiple pairwise comparisons between WT and KO mice at both ages. For amplitude-amplitude coupling, Fisher Z-transformed values of spearman’s correlation were averages and compared between KO and WT mice at both age using a two-way ANOVA with repeated measures followed by post hoc multiple pairwise comparisons for all six correlations including theta-low gamma, theta-high gamma, low alpha-low gamma, low alpha-high gamma, high alpha-low gamma, and high alpha-high gamma.

## Results

### Absolute alpha, beta, gamma and total EEG power was increased in KO mice compared to WT controls. Gamma power increased when both the WT and KO animals developed from juveniles to adults

Power spectra indicated that EEG power was increased in KO mice compared to WT controls at both P40 and P70 in the frequency range of alpha, beta, and gamma (8-100Hz) in home cage **(Figure 2A)**, light-dark arena **(Figure 2B)** and open field arena **(Figure 2C)**. In home cage recording **(Figure 2D)**, we observed that absolute EEG power in alpha, beta, and gamma, including low and high gamma frequency bands, was increased in KO mice compared to WT controls at both P40 and P70. Alongside, total power was also increased in the KO mice at both ages. However, absolute EEG power in delta and theta bands was similar between KO and WT mice at both ages. Regarding developmental changes, absolute gamma power was increased in both WT and KO mice from P40 to P70.

**Figure 2:**
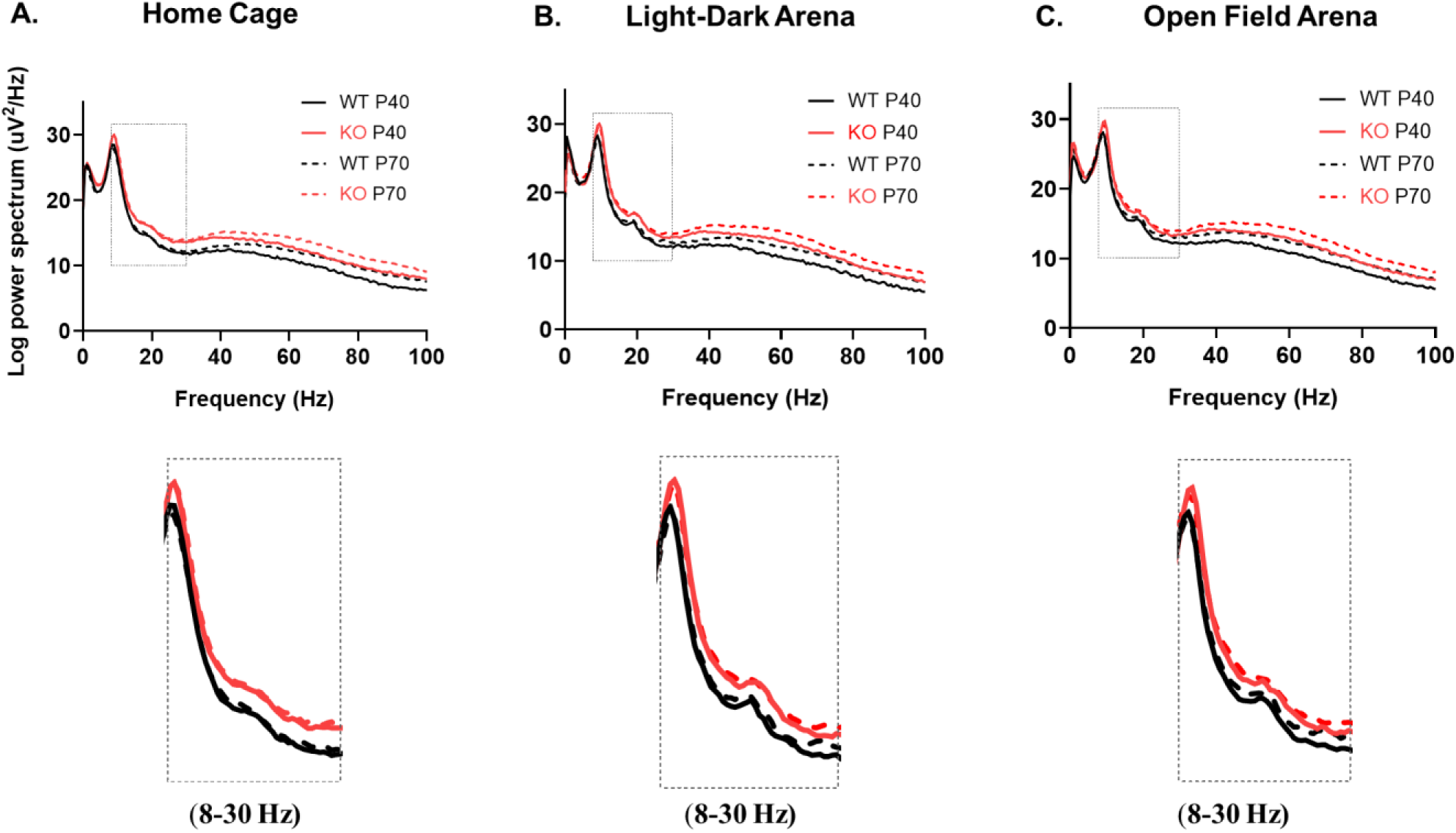
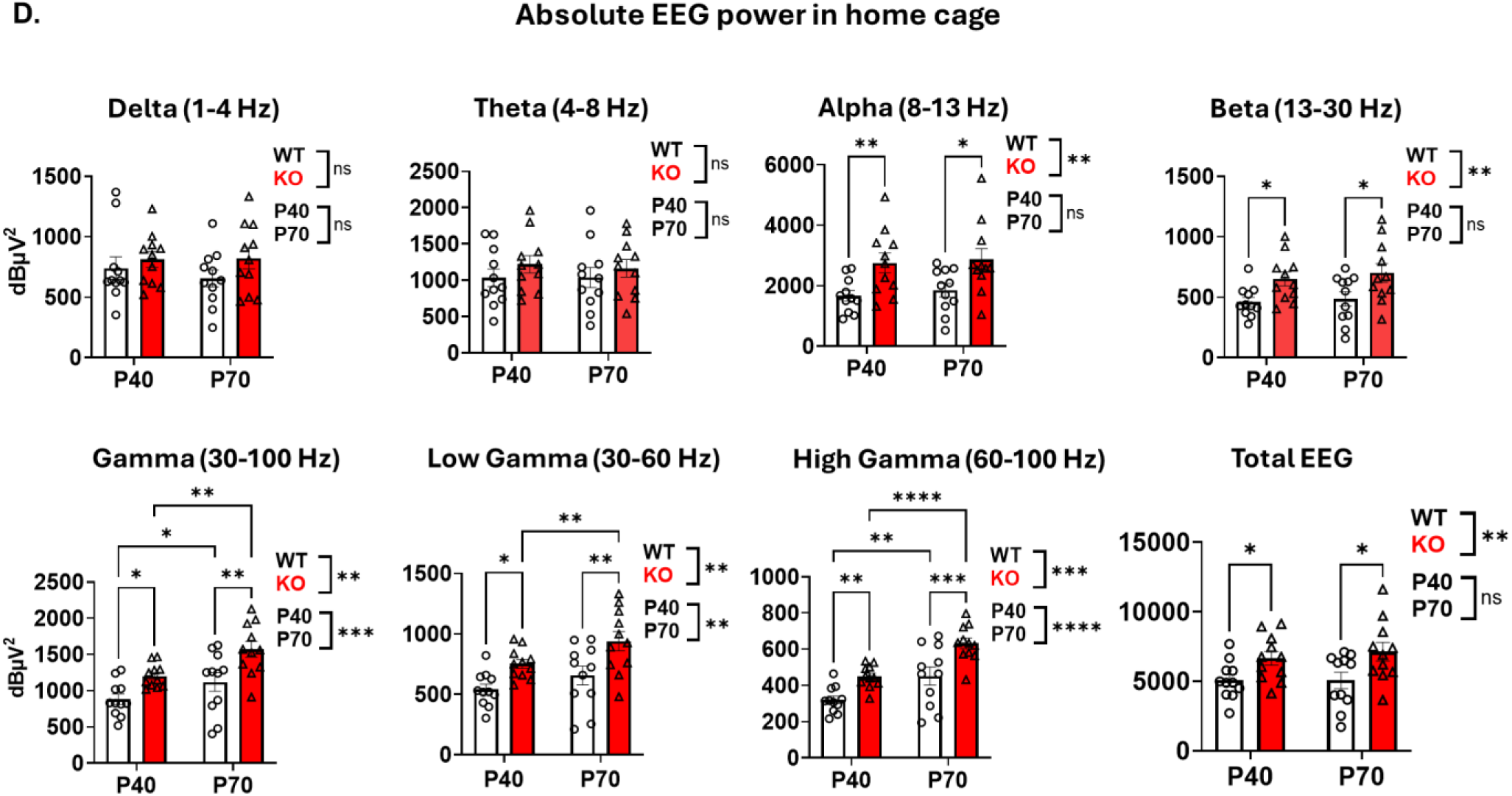
Alpha, beta, gamma and total EEG power was increased in KO compared to WT mice at both ages, and gamma power increased from P40 to P70. (A, B, & C) Power spectra of KO and WT mice at both ages in the home cage, light-dark, and open field arenas. (D) Comparison of absolute EEG power between WT and KO, in delta, theta, alpha, beta, and gamma, low and high gamma frequency band, and total power in home cage recordings at both ages. No significant interaction (Genotype X Age) was found. Data represent mean ± SEM for KO (n=11) and WT (n=11) mice. Statistical significance was assessed using two-way repeated measures ANOVA, followed by post hoc multiple pairwise comparisons. *: P < 0.05, **: P < 0.01, ***: P < 0.001, ****: P < 0.0001. Detailed descriptions of statistical analysis methods are presented in Methods.

Overall, similar increases in alpha, beta, gamma, including low and high gamma bands, and total EEG power were found in light-dark and open field arena recordings at both ages, as summarized in **Table 1 & 2**. Therefore, it is evident that EEG power consistently increased in the KO mice in the frequency range of 8-100Hz, compared to WT controls at juvenile and adult ages, and gamma power was increased from juvenile to adult ages in both genotypes, regardless of the three recording conditions used.

**Table 1:**
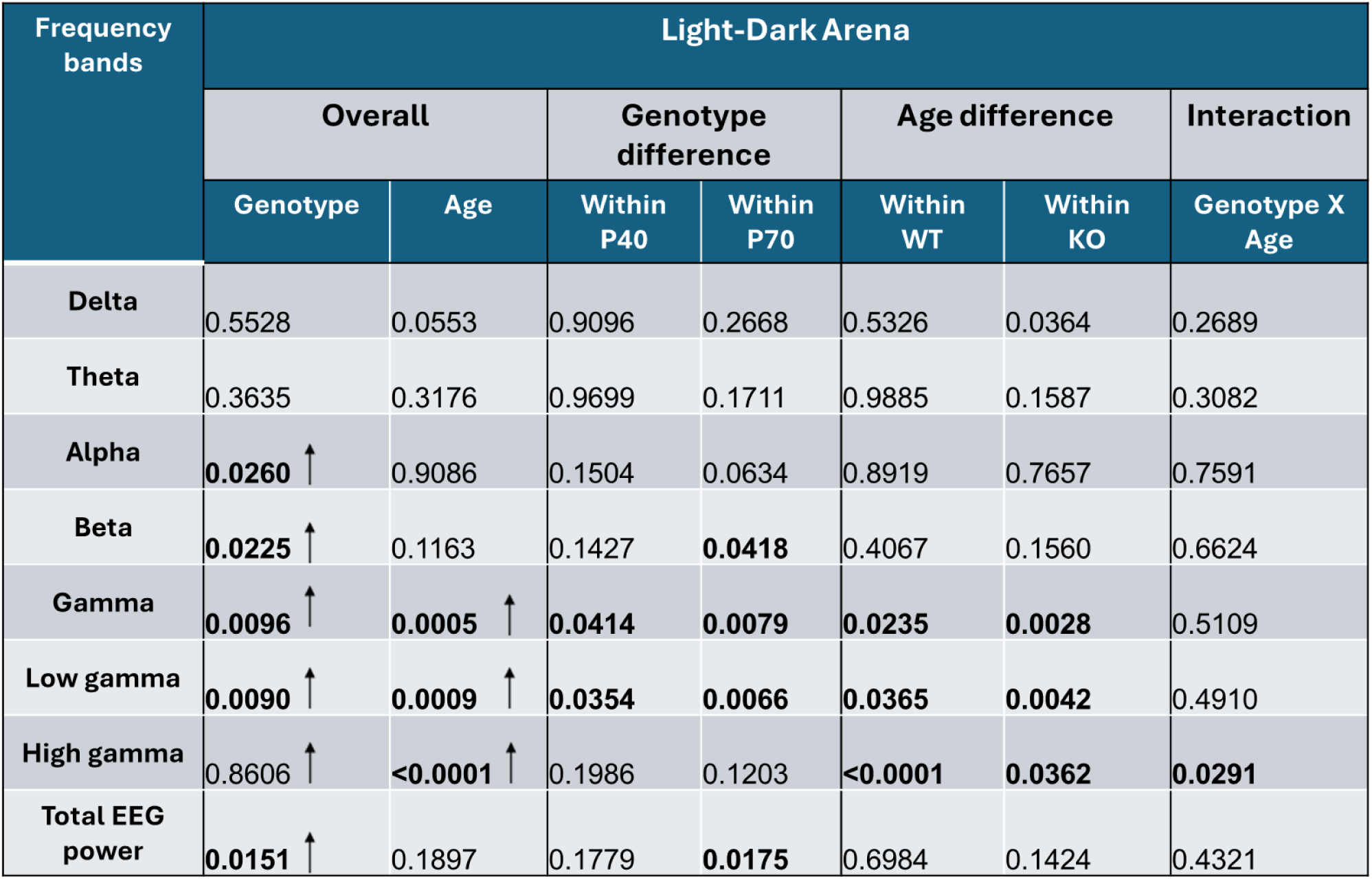
P-value summary based on Two-way ANOVA repeated measures comparing absolute EEG power in light-dark arena recording between KO and WT mice. Summary of the P-values from two-way repeated measures ANOVA comparing absolute EEG power in KO and WT mice at P40 and P70, in light-dark arena. Arrowhead and bold text indicate significant changes (increase) in power between genotypes or ages.

| Frequency bands | Light-Dark Arena |  |  |  |  |  |  |
| --- | --- | --- | --- | --- | --- | --- | --- |
|  | Overall |  | Genotype difference |  | Age difference |  | Interaction |
|  | Genotype | Age | Within P40 | Within P70 | Within WT | Within KO | Genotype X Age |
| Delta | 0.5528 | 0.0553 | 0.9096 | 0.2668 | 0.5326 | 0.0364 | 0.2689 |
| Theta | 0.3635 | 0.3176 | 0.9699 | 0.1711 | 0.9885 | 0.1587 | 0.3082 |
| Alpha | **0.0260** ↑ | 0.9086 | 0.1504 | 0.0634 | 0.8919 | 0.7657 | 0.7591 |
| Beta | **0.0225** ↑ | 0.1163 | 0.1427 | **0.0418** | 0.4067 | 0.1560 | 0.6624 |
| Gamma | **0.0096** ↑ | **0.0005** ↑ | **0.0414** | **0.0079** | **0.0235** | **0.0028** | 0.5109 |
| Low gamma | **0.0090** ↑ | **0.0009** ↑ | **0.0354** | **0.0066** | **0.0365** | **0.0042** | 0.4910 |
| High gamma | 0.8606 ↑ | **<0.0001** ↑ | 0.1986 | 0.1203 | **<0.0001** | **0.0362** | **0.0291** |
| Total EEG power | **0.0151** ↑ | 0.1897 | 0.1779 | **0.0175** | 0.6984 | 0.1424 | 0.4321 |

### Absolute EEG power was stable across recording conditions

We observed that absolute EEG power was stable across different time points when recorded from the home cage and followed by either the light-dark or the open field arenas, for all frequency bands analyzed (delta, theta, alpha, gamma [low and high gamma], and total EEG power) **(Figure 3A, B)**.

**Figure 3:**
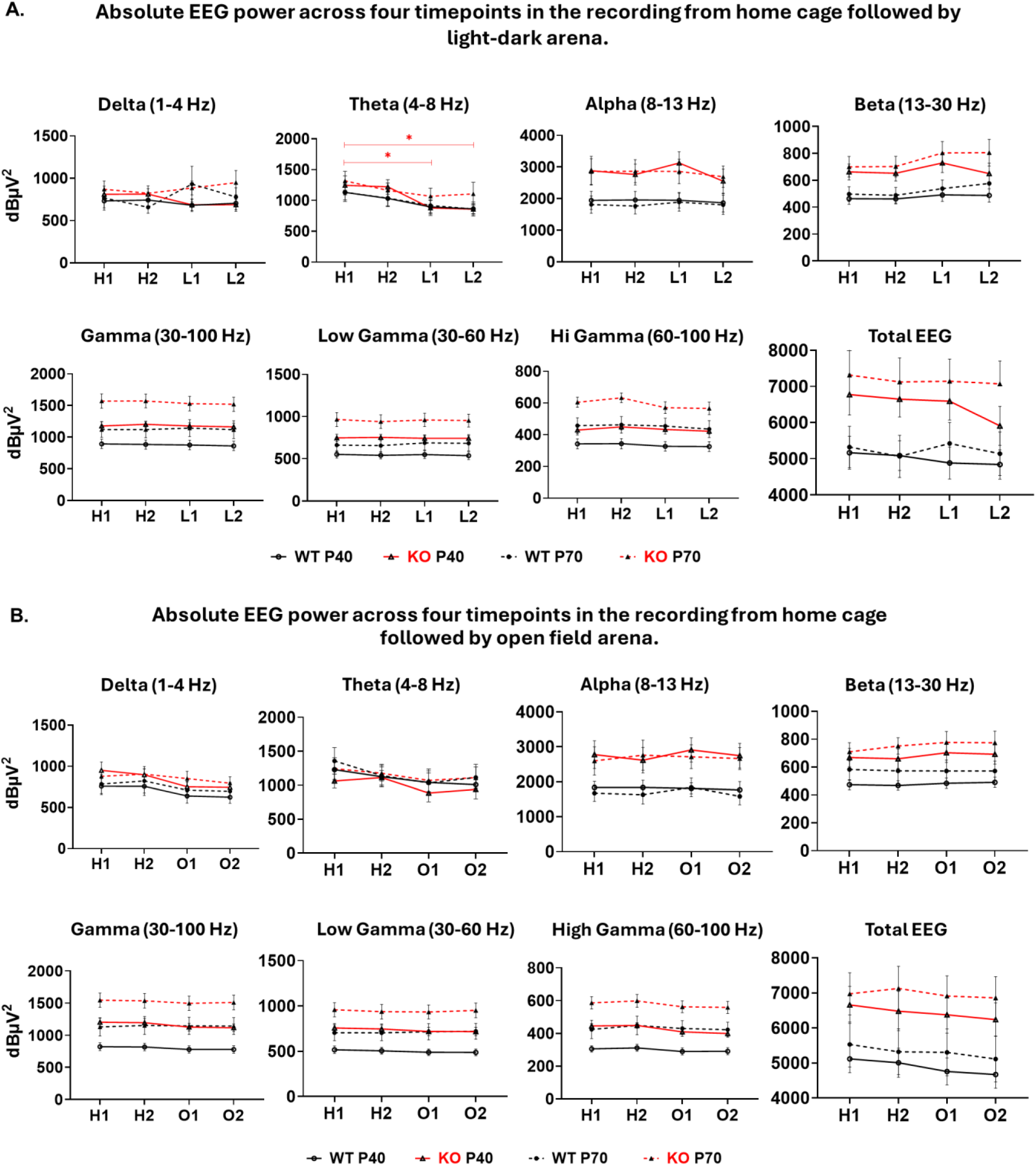
Absolute EEG power was stable across different recording settings in WT and KO mice at both P40 and P70. (A) Comparison of absolute EEG power at H1 (15–20 min) and H2 (25–30 min) in the home cage and L1 (15–20 min) and L2 (25–30 min) in the light-dark arena. (B) Comparison of absolute EEG power at H1, H2, and O1 (15–20 min) and O2 (25–30 min) in the open field arena. Data represent mean ± SEM for KO (n=11) and WT (n=11) mice. Statistical significance was assessed using three-way repeated measures ANOVA, followed by post hoc multiple pairwise comparisons. *P < 0.05. Detailed statistical methods are described in the Methods section.

One exception was that in KO mice at P40, theta power significantly decreased from H1 (home cage recording, 15–20 minutes) to L1 (light-dark arena recording, 15–20 minutes) and from H1 to L2 (light-dark arena recording, 25–30 minutes) **(Figure 3A)**. These reductions in theta power were not observed in KO mice at P70 or WT mice **(Figure 3A)**, or in recordings from home cage and followed by the open field arena **(Figure 3B)**. No changes in other frequency bands were observed.

### Peak alpha frequency was increased in KO mice compared to WT controls at both ages

Peak alpha frequency, defined as the frequency at which the maximum power occurs within the alpha band, shifts from lower to higher frequencies during brain maturation (24). Notably, peak alpha frequency has been found to be altered in several neurodevelopmental disorders, including attention deficit hyperactivity disorder (ADHD), ASD, and FXS (19, 36, 37). In current study, peak alpha frequency was increased in KO mice compared to WT controls in recordings from light-dark and open field arenas, at P40 and P70 **(Figure 4, A & B)**. Interestingly, it did not change in home cage recording between WT and KO mice. There was no developmental change between P40 and P70.

**Figure 4:**
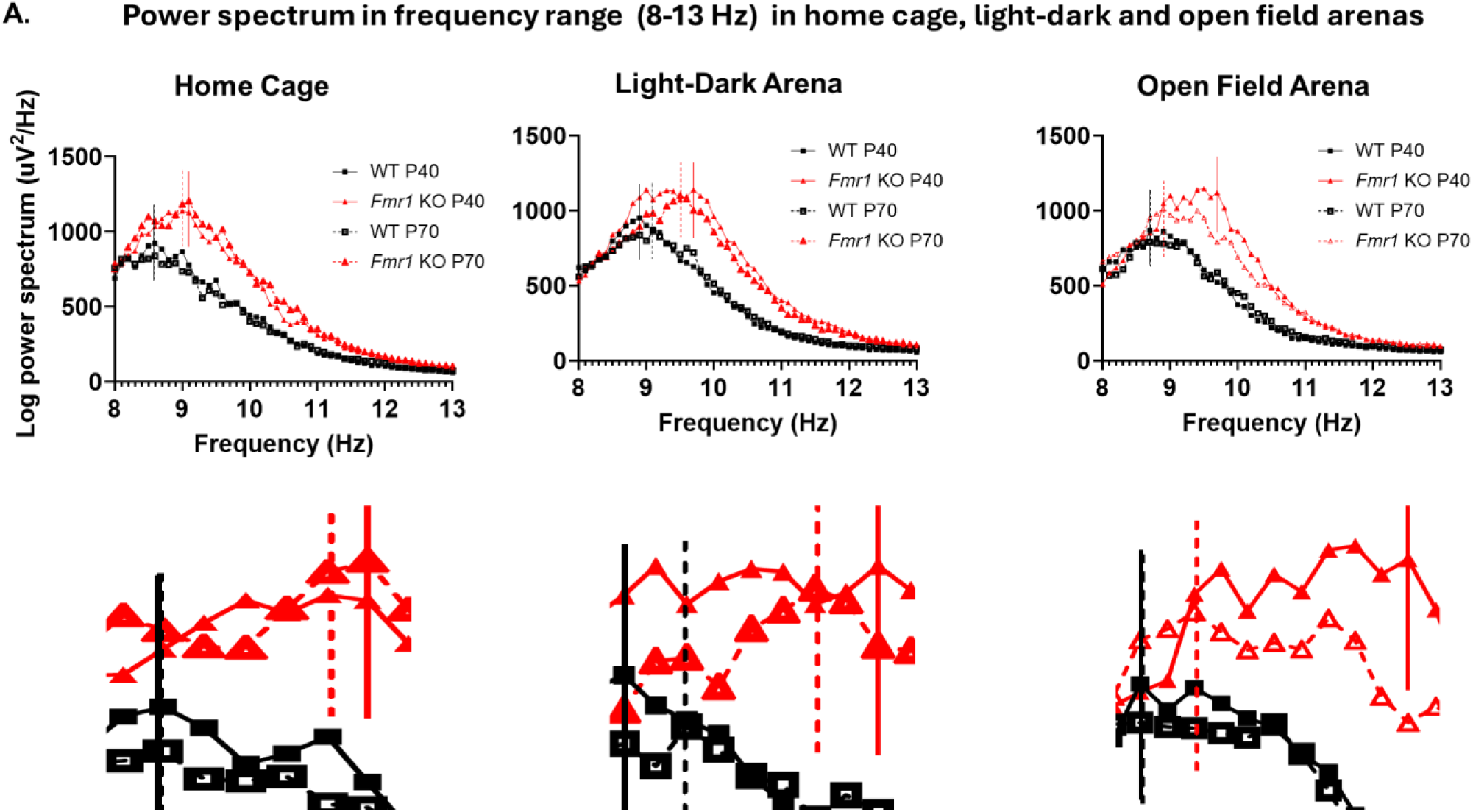
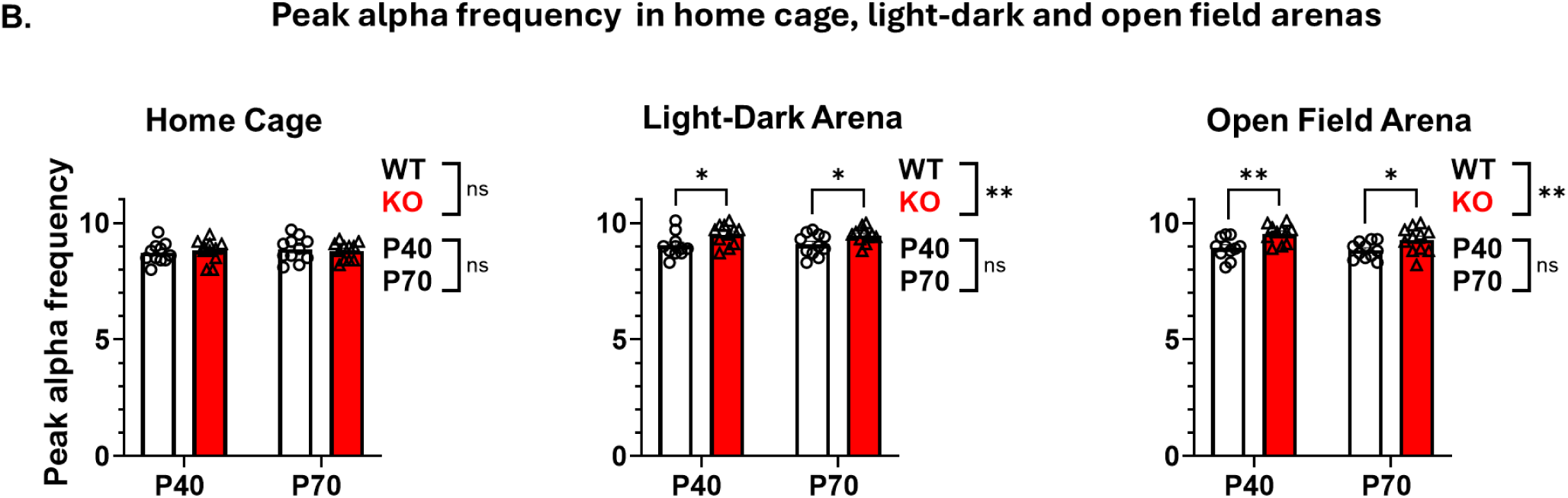
Peak alpha frequency was increased in KO mice at P40 and P70 in the light-dark and open field arenas. (A) Average log power spectrum in the alpha frequency range (8-13 Hz). Solid and dotted lines (orange and black) point out the peak alpha frequency for all 4 groups. (B) Comparison of peak alpha frequency between WT and KO mice at P40 and P70 in the home cage, light-dark, and open field arenas. No significant interaction (Genotype X Age) was found. Data represent mean ± SEM for KO (n=11) and WT (n=11) mice. Statistical significance was assessed using two-way repeated measures ANOVA, followed by post hoc multiple pairwise comparisons. *P < 0.05, **P < 0.01. Detailed statistical methods are described in the Methods section.

### Theta-beta ratio decreases in KO mice at both ages compared to WT controls

The theta-beta power ratio is one of the most well-established EEG biomarkers for ADHD (38, 39), where it is characterized by elevated slow theta waves and reduced fast beta waves. Given prior evidence of increased theta power in FXS (11, 40), it is plausible that this marker may also be affected in this condition. Theta-beta ratio was decreased in KO mice as compared to WT controls across all recording conditions, except in the light-dark arena at P70. No developmental difference (P40 vs P70) was found in theta-beta ratio **(Figure 5)**.

**Figure 5:**
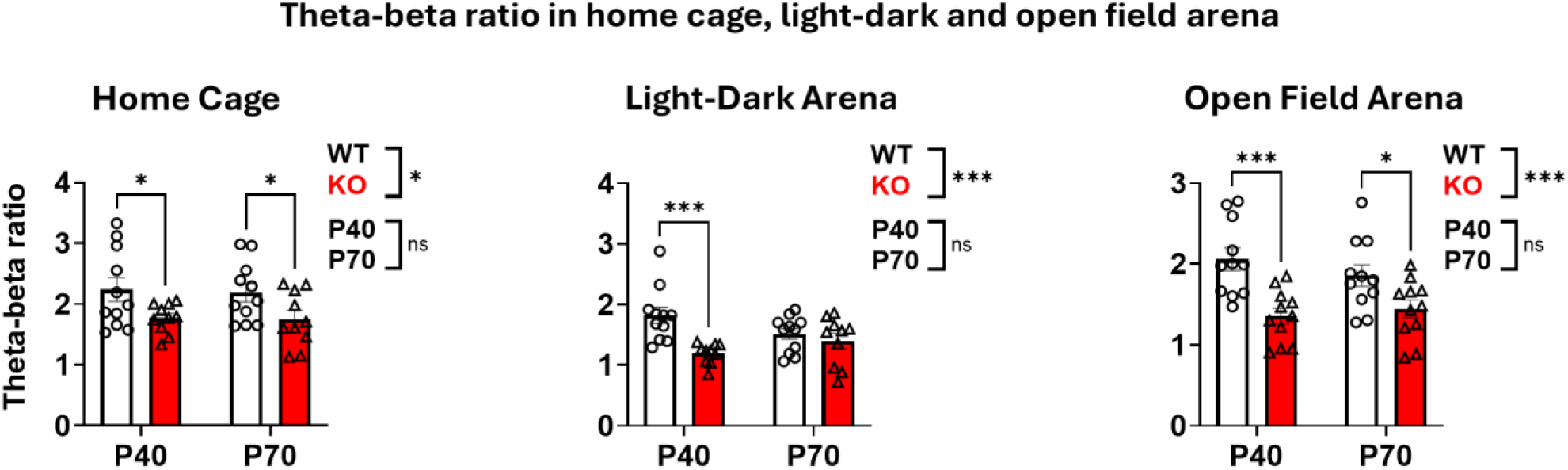
Theta-beta ratio decreased in KO compared to WT mice at both ages, except in the light-dark arena at P70. Comparison of theta-beta ratio between WT and KO mice at P40 and P70 in home cage, light-dark and open field arenas. No significant interaction (Genotype X Age) was found. Data represent mean ± SEM for KO (n=11) and WT (n=11) mice. Statistical significance was assessed using two-way repeated measures ANOVA, followed by post hoc multiple pairwise. *P < 0.05, **P < 0.01, ***P < 0.001. Detailed statistical methods are described in the Methods section.

### Relative theta EEG power decreased in KO mice compared to WT controls. Relative gamma power increased from the juvenile to adult stage

In home cage recording, relative EEG theta power decreased in KO mice compared to WT controls at P70; however, no change was observed at P40. Alongside, no genotype difference was found in relative delta, alpha, beta, and gamma (including both low and high gamma) EEG power at both ages **(Figure 6)**. Relative gamma power (including low and high gamma) increased from juvenile to adult ages in both WT and KO mice.

**Figure 6:**
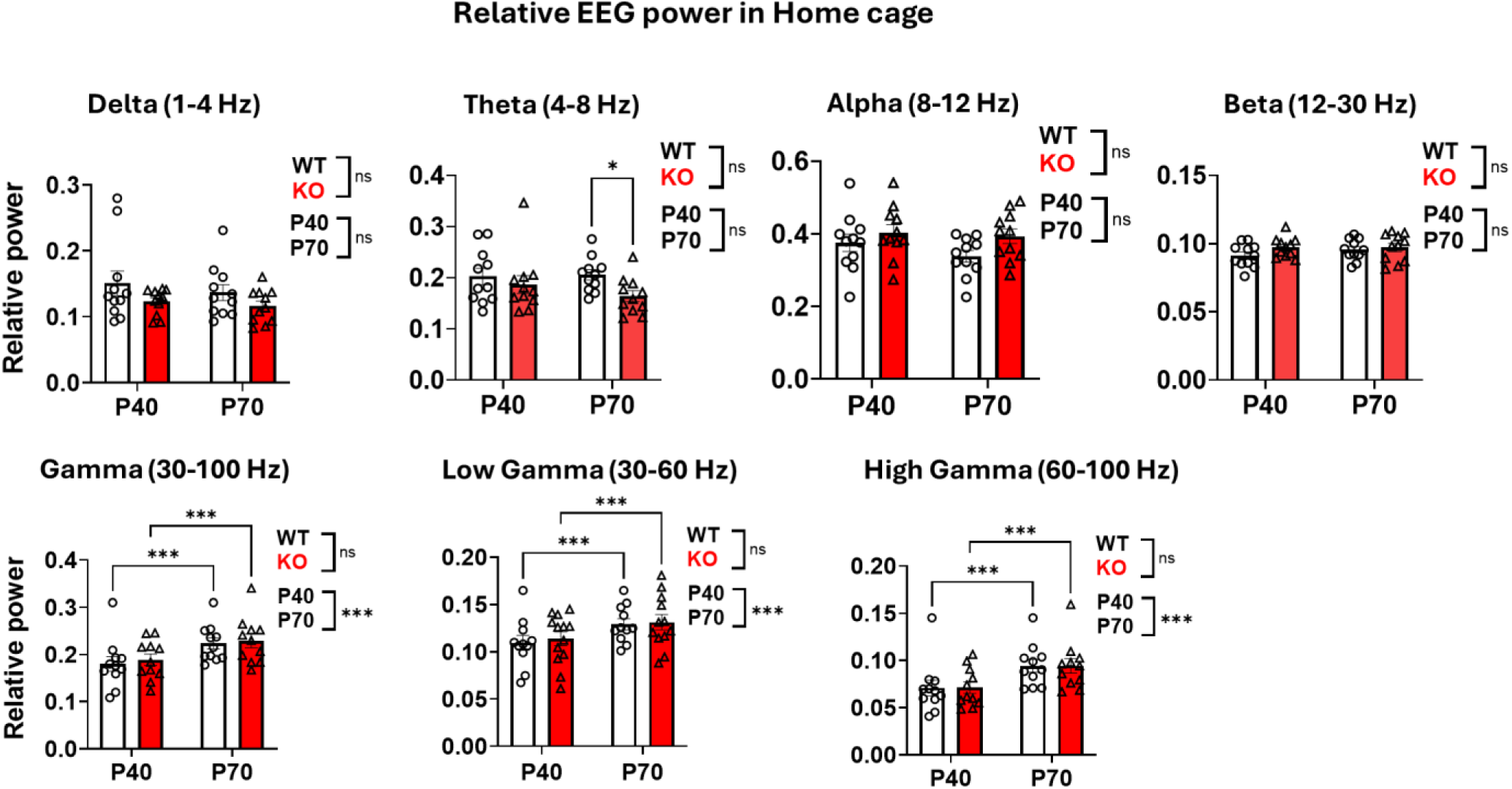
Relative theta power decreased in KO mice at P70; relative gamma power increased from P40 to P70. Figure shows relative EEG power for delta, theta, alpha, beta, and gamma, including low and high gamma, in home cage recording. No significant interaction (Genotype X Age) was found. Data represent mean ± SEM for KO (n=11) and WT (n=11) mice. Statistical significance was assessed using two-way repeated measures ANOVA, followed by post hoc multiple pairwise comparisons. *P < 0.05, **P < 0.01, ***P < 0.001. Detailed statistical methods are described in the Methods section.

In the light-dark arena, no genotypic difference was found in all frequency bands **(Table 3)**. However, high gamma power displayed developmental changes, driven by an increase from P40 to P70 in WT controls.

**Table 2:** P-value summary based on Two-way ANOVA repeated measures comparing absolute EEG power in open field arena recording between KO and WT mice. Summary of the P-values from two-way repeated measures ANOVA comparing absolute EEG power in KO and WT mice at P40 and P70, in open field arena. Arrowhead and bold text indicate significant changes (increase) in power between genotypes and ages.

| Frequency bands | Open Field Arena |  |  |  |  |  |  |
| --- | --- | --- | --- | --- | --- | --- | --- |
|  | Overall |  | Genotype difference |  | Age difference |  | Interaction |
|  | Genotype | Age | Within P40 | Within P70 | Within WT | Within KO | Genotype X Age |
| Delta | 0.3097 | 0.1815 | 0.3519 | 0.3749 | 0.3337 | 0.3456 | 0.9626 |
| Theta | 0.4838 | 0.1500 | 0.8069 | 0.4048 | 0.1722 | 0.4821 | 0.6499 |
| Alpha | **0.0060** ↑ | 0.7535 | **0.0211** | **0.0214** | 0.8268 | 0.8216 | 0.9962 |
| Beta | **0.0193** ↑ | 0.1400 | **0.0419** | **0.0401** | 0.2955 | 0.2848 | 0.9862 |
| Gamma | **0.0074** ↑ | **<0.0001** ↑ | **0.0170** | **0.0103** | **0.0010** | **0.0005** | 0.8386 |
| Low gamma | **0.0116** ↑ | **<0.0001** ↑ | **0.0188** | **0.0188** | **0.0014** | **0.0014** | 0.9997 |
| High gamma | **0.0057** ↑ | **<0.0001** ↑ | **0.0238** | **0.0053** | **0.0008** | **0.0001** | 0.5673 |
| Total EEG power | **0.0122** ↑ | 0.2839 | **0.0479** | **0.0288** | 0.5233 | 0.3748 | 0.8570 |

**Table 3:**
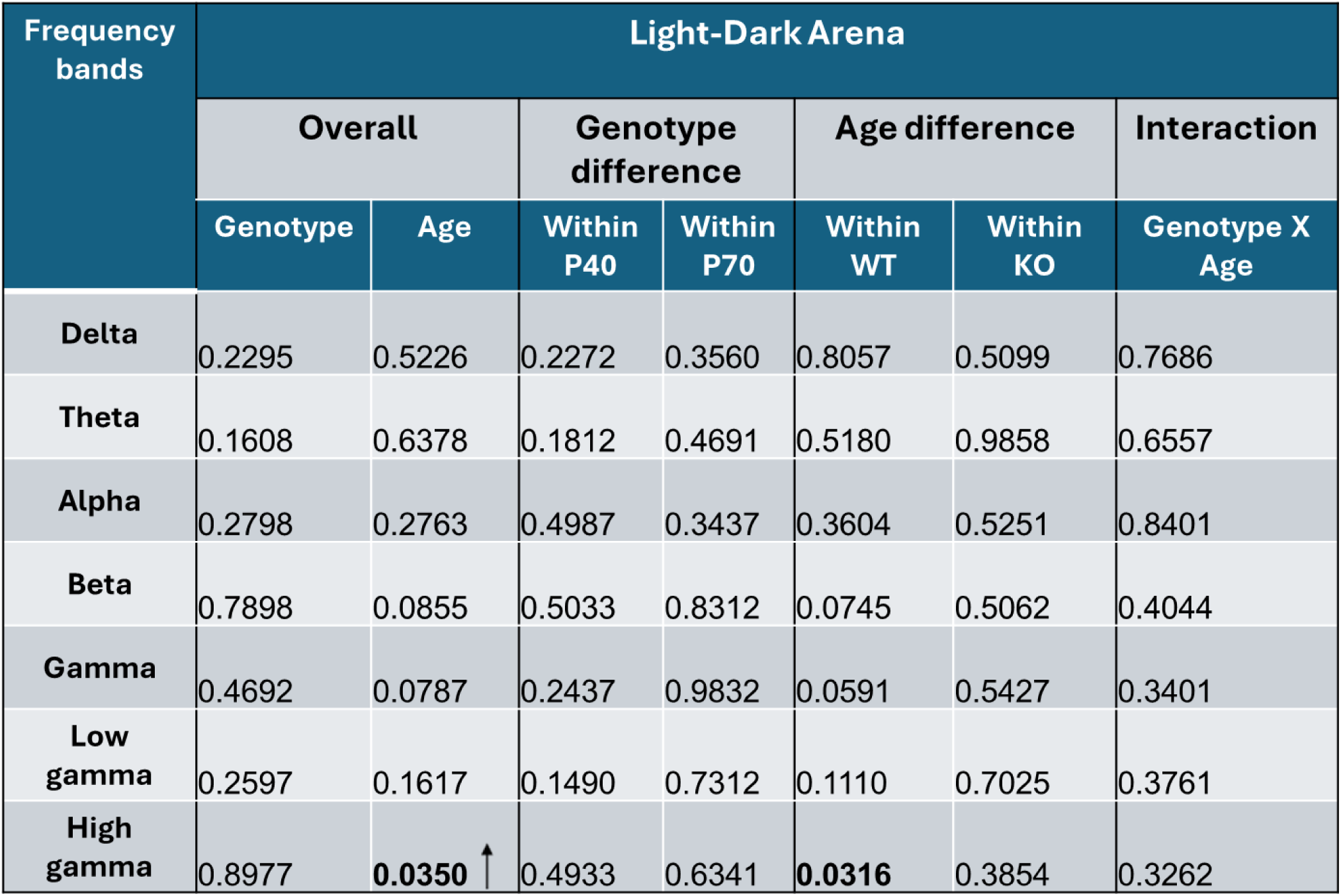
P-value summary based on Two-way ANOVA repeated measures comparing relative EEG power in light-dark arena recording between KO and WT mice. Summary of the P-values from two-way repeated measures ANOVA comparing relative EEG power in KO and WT mice at P40 and P70, in the light-dark arena. Arrowhead and bold text indicate a significant increase in relative power within WT mice from P40 to P70.

In the open field arena, relative theta power decreases in KO mice compared to WT controls at both P40 and P70 **(Table 4)**. Alongside, relative alpha power was increased in KO mice compared to WT controls. No other genotype difference was found in relative delta, beta, and gamma (low and high gamma) power at both ages. Regarding developmental changes, relative alpha power decreased from P40 to P70 (**Table 4**). Moreover, relative gamma power (including low and high gamma) increased from P40 to P70.

**Table 4:** P-value summary based on Two-way ANOVA repeated measures comparing relative EEG power in open field arena recording between KO and WT mice. Summary of the P-values from two-way repeated measures ANOVA comparing relative EEG power in KO and WT mice at P40 and P70, in the open field arena. Arrowhead and bold text indicate a significant increase or decrease in relative power for genotype and age differences.

| Frequency bands | Open Field Arena |  |  |  |  |  |  |
| --- | --- | --- | --- | --- | --- | --- | --- |
|  | Overall |  | Genotype difference |  | Age difference |  | Interaction |
|  | Genotype | Age | Within P40 | Within P70 | Within WT | Within KO | Genotype X Age |
| Delta | 0.3947 | 0.6402 | 0.6778 | 0.2903 | 0.9120 | 0.4427 | 0.5340 |
| Theta | **0.0010** ↓ | 0.8285 | **0.0010** | **0.0180** | 0.6500 | 0.4497 | 0.3941 |
| Alpha | **0.0247** ↑ | **0.0055** ↓ | 0.0739 | **0.0255** | **0.0198** | 0.0761 | 0.6445 |
| Beta | 0.8387 | 0.0674 | 0.5852 | 0.8493 | 0.0720 | 0.4132 | 0.4605 |
| Gamma | 0.8265 | **0.0002** ↑ | 0.4544 | 0.6900 | **0.0009** | **0.0194** | 0.3503 |
| Low gamma | 0.6982 | **0.0002** ↑ | 0.3383 | 0.7244 | **0.0008** | **0.0244** | 0.3019 |
| High gamma | 0.9924 | **0.0004** ↑ | 0.6735 | 0.6851 | **0.0019** | **0.0210** | 0.4582 |

### Increased phase-amplitude cross frequency coupling in KO mice compared to WT controls, and in adults compared to juveniles

Phase amplitude coupling assesses how the amplitude of high-frequency activity is modulated by the phase of low-frequency oscillations. In the brain, strong coupling functions as a clocking mechanism to produce perceptual windows that, in turn, integrate and separate temporally relevant and irrelevant information (41). Interestingly, one study with FXS patients reported no group differences between FXS patients and neurotypical controls for phase amplitude coupling (42). However, there is not much available in the literature for this measurement in FXS rodent models. In the current study, the strength of phase amplitude coupling, measured between the amplitude of gamma band (30–100 Hz) and the phase of lower frequency oscillations (4–12 Hz), was higher in KO mice compared to WT controls in home cage and the light-dark arena, but did not reach statistical significance in the open field arena **(Figure 7, A and B)**. Regarding developmental changes, PAC value increased from P40 to P70. The difference reached statistical significance in the recordings from the light-dark and open field arenas **(Figure 7, A and B)**.

**Figure 7:**
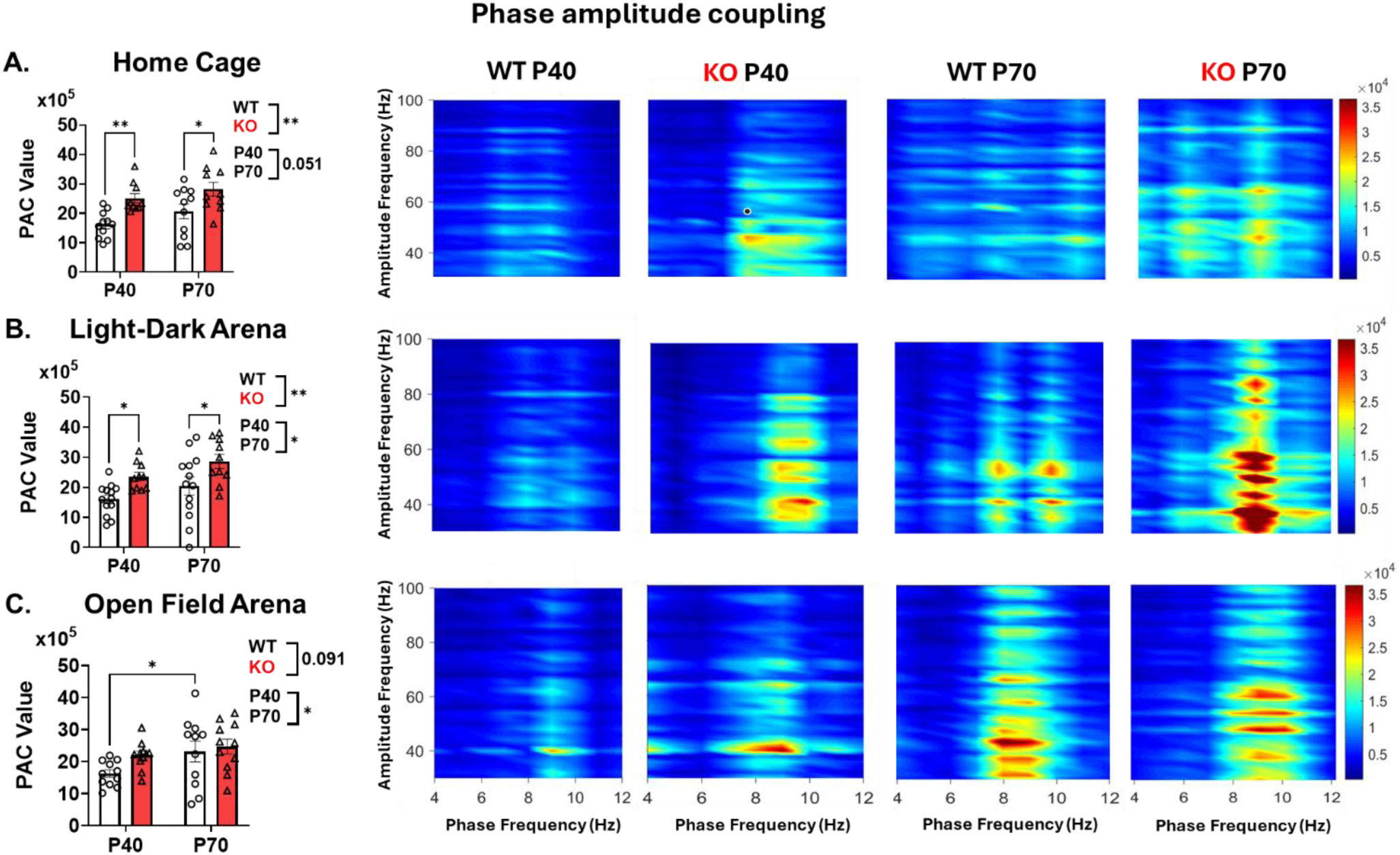
Phase-amplitude cross frequency coupling was generally increased in KO mice compared to WT controls, and in adults compared to juveniles. (A, B, & C) Shows the quantification of phase-amplitude coupling and phase-amplitude map for recordings in home cage, light-dark, and open field arenas. The left panels show the quantification of total mean vector length (MVL) values and comparison between WT and KO mice at P40 and P70. The right panels show representative phase-amplitude heat map between the amplitude of the gamma band (30 – 100 Hz, amplitude-frequency) and the phase of the low-frequency band (4 – 12 Hz, phase frequency). The color scale is representative of MVL values, a measurement of coupling strength. Data represent mean ± SEM for KO (n=11) and WT (n=11) mice. Statistical significance was assessed using two-way repeated measures ANOVA, followed by post hoc analysis. *P < 0.05, **P < 0.01. Detailed statistical methods are described in the Methods section.

### Negative theta-high gamma and positive high alpha-high gamma amplitude-amplitude cross frequency coupling were increased in KO mice

Amplitude-amplitude coupling particularly assesses the associations of alpha and theta activity with gamma activity over time (11). Although less frequently studied than phase-amplitude coupling, amplitude-amplitude coupling has been explored in FXS, particularly during resting-state EEG. Studies have identified altered theta-gamma amplitude-amplitude coupling in FXS, characterized by an inverse relationship between increased theta power and decreased gamma power (11, 42). In the current study, we investigated the six different sets of amplitude-amplitude coupling, between theta-low gamma, theta-high gamma, low alpha-low gamma, low alpha-high gamma, high alpha-low gamma, and high alpha-high gamma. All six of these amplitude – amplitude coupling were calculated from EEG recordings of both WT and KO mice at both P40 and P70 in home cage, light-dark and open field arenas.

Consistent changes that we observed included that compared to WT controls, KO mice had greater magnitudes of negative theta-high gamma amplitude – amplitude coupling, as well as greater magnitudes of positive high alpha-high gamma amplitude – amplitude coupling, at P40 in home cage recording, and at both P40 and P70 in the light-dark arena **(Figure 8, A and B)**.

**Figure 8:**
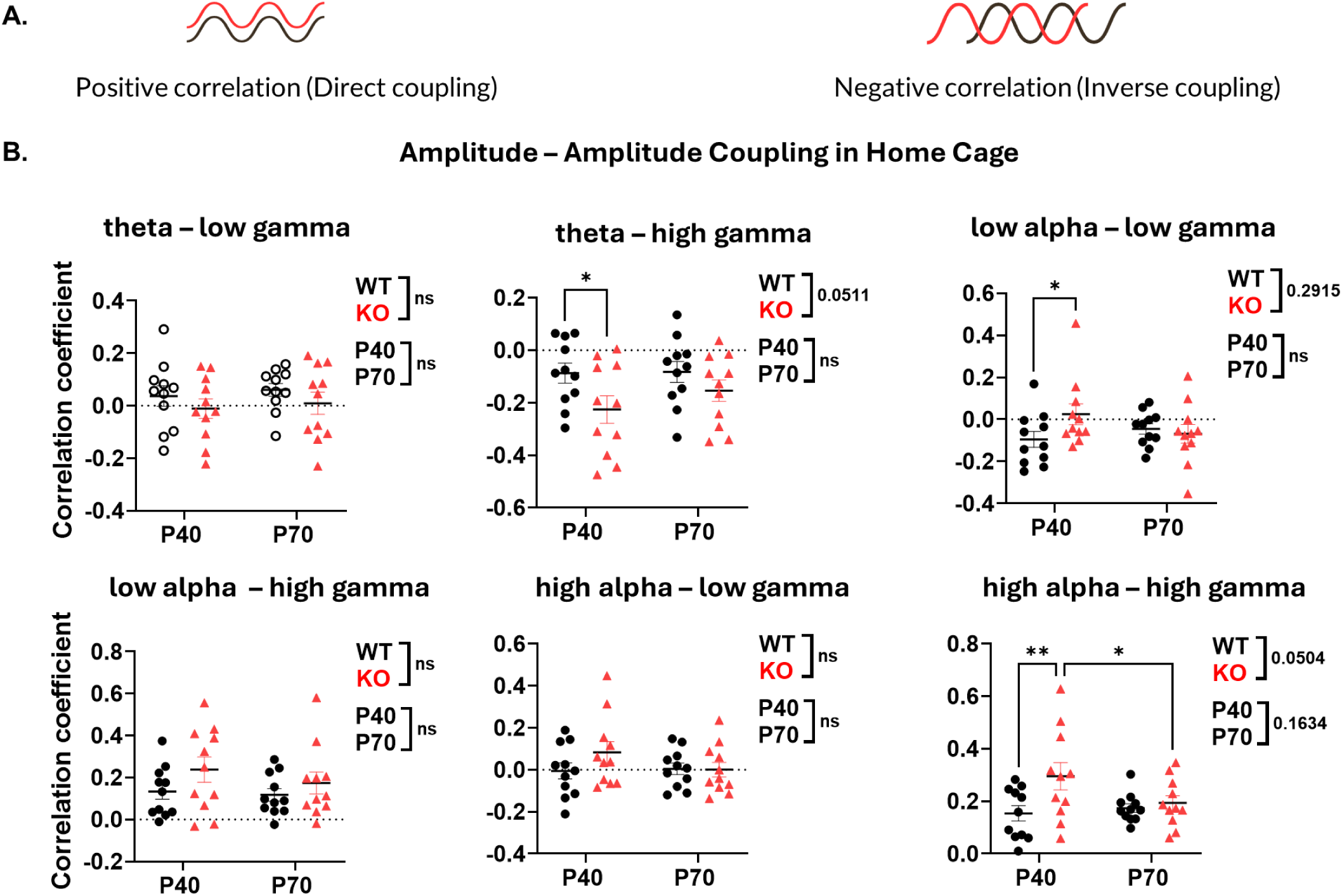
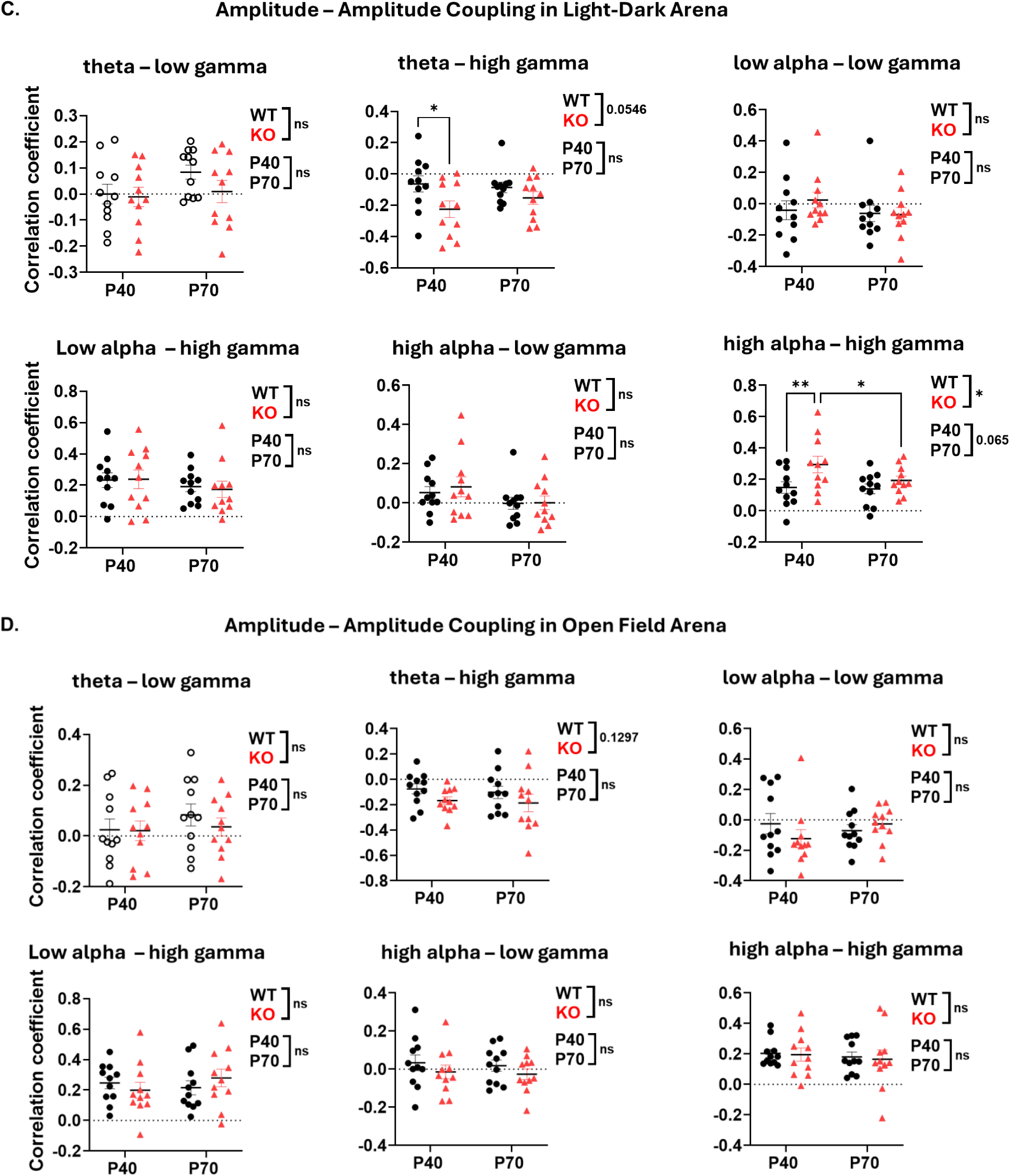
Amplitude-amplitude cross frequency coupling between low frequency (theta and alpha) and high frequency power (gamma). (A) Visual representation of positive (direct) and negative (inverse) coupling between two waves. (B, C & D) Cross frequency amplitude coupling including theta-low gamma, theta-high gamma, low alpha-low gamma, low alpha-high gamma, high alpha-low gamma, and high alpha-high gamma is compared between WT and KO mice at P40 and P70 in home cage, light-dark and open field arenas. Data represent mean ± SEM for KO (n=11) and WT (n=11) mice. Statistical significance was assessed using two-way repeated measures ANOVA, followed by post hoc analysis. *P < 0.05, **P < 0.01. Detailed statistical methods are described in the Methods section.

No genotype difference was found in any of the six couplings in the open field arena (Figure 8C).

### Multiscale entropy was unchanged between KO and WT mice at both ages

Finally, EEG signal complexity is regarded as an indicator of brain maturation and cognitive functioning, as it has been shown to increase with age (43). Moreover, its increase has been shown to be sensitive to the maturation patterns of specific sensory brain regions (44). Although findings are inconsistent, several studies on ADHD and ASD patients have generally reported a reduction in EEG signal complexity compared to the controls (45-47), while another study found that individuals with ADHD exhibit reduced EEG signal complexity specifically within the alpha frequency band (48). Multiscale entropy is a robust technique for quantitatively assessing signal complexity, as it investigates temporal complexity of the signal at multiple time scales (24). One study revealed that the signal complexity in FXS participants was reduced compared to the controls (24). Building on these findings, we evaluated signal complexity in our current study. **Figure 9A** illustrates multi-scale entropy (MSE) at timescale 1-40 in WT and KO at both P40 and P70. Complexity index, which is the area under the curve for MSE 1-40, showed no difference between WT and KO mice at both P40 and P70 in all three recording conditions **(Figure 9B)**. Furthermore, MSE across lower timescales (1-20) and higher timescale (21-40) also remained unchanged between WT and KO mice at both ages, across all three conditions **(Figure 9C).**

**Figure 9:**
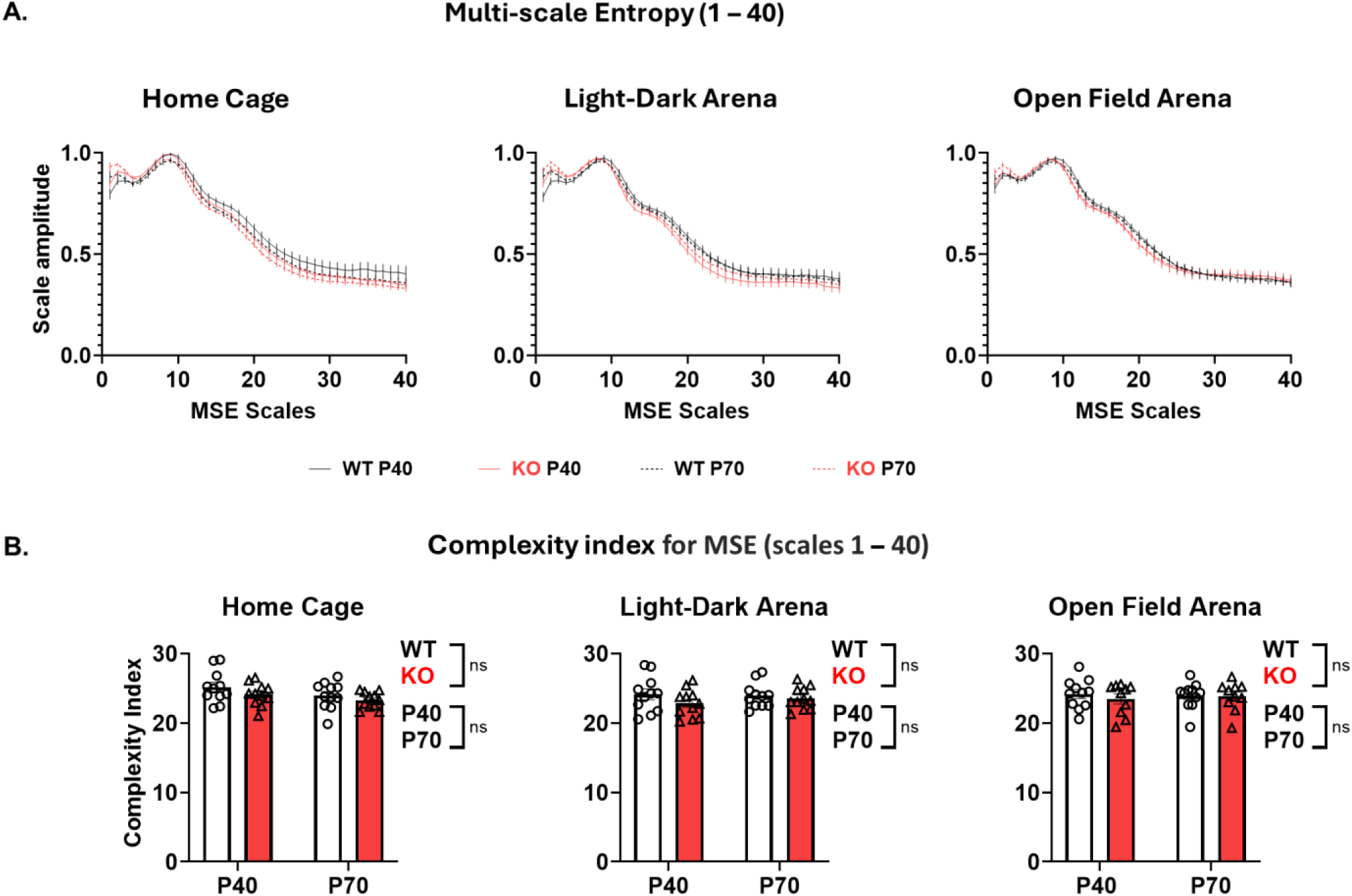
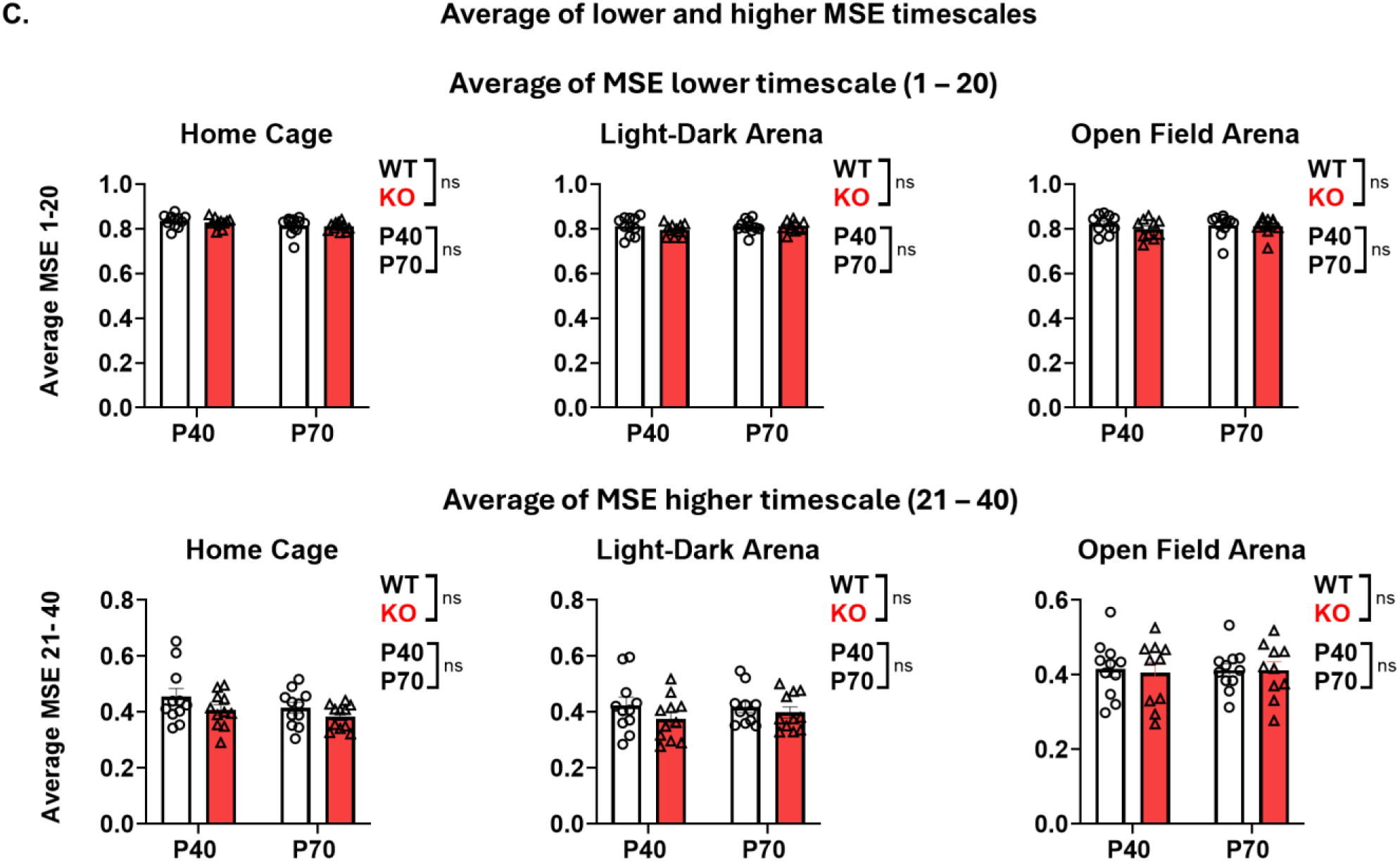
No change in MSE between WT and KO mice at P40 and P70. (A) Average amplitude of MSE at timescales (1-40) for WT and KO mice at P40 and P70. (B) Complexity index (area under the curve) for MSE (1-40) was compared between WT and KO mice at P40 and P70 in home cage, light-dark and open field arenas. (C) Average of MSE lower timescale (MSE 1-20) and higher timescale (MSE 21-40), compared between WT and KO mice at P40 and P70 in home cage, light-dark and open field arenas. Data represent mean ± SEM for KO (n=11) and WT (n=11) mice. Statistical significance was assessed using two-way repeated measures ANOVA. Detailed statistical methods are described in the Methods section.

## Discussion

### EEG as a translational biomarker for FXS

As we know, currently no interventions exist for FXS, and many drugs have failed in numerous clinical trials. Thus, there is a need for an effective translational biomarker (6-9). EEG has been suggested as such a biomarker for FXS, with potential indicators identified in the EEG alterations observed in FXS patients. These include elevated gamma power at rest, reduced gamma phase-locking in sensory cortices, and increased amplitudes of event-related potentials. Such deviations could signal cortical hyperactivity, which may result from a disruption in the balance between excitatory systems, driven by glutamate, and inhibitory systems, mediated by GABA, in FXS (7).

Although the exact underlying mechanisms generating gamma oscillation are still being debated, the inhibitory GABAergic interneurons have been proposed to play a critical role in synchronizing cortical activities at gamma frequencies (49-51). Additionally, dysfunctional inhibitory neurons and associated circuits have been identified in the rodent models of Fragile X Syndrome, which lead to impaired inhibitory control of cortical synchronization (52-54).

There is some evidence from clinical trials showing that treatment-related changes in EEG could be associated with clinical improvements for various neurodevelopmental conditions including ASD (6). However, these findings require replication and extension to other medications. While the use of EEG characteristics as biomarkers is still in the early phases, the current research is promising and could signal the potential emergence of an effective translational biomarker (6).

### Cross frequency coupling in FXS mouse models

Cross-frequency coupling (CFC) has been developed as a method to assess moment-to-moment interaction between oscillations of different frequencies within EEG signals (55). Two primary forms of CFC, Phase-Amplitude Coupling and Amplitude-Amplitude Coupling offers insights into neural dynamics and network integration (55).

A recent EEG study in children aged 6–11 with autism revealed heightened alpha-gamma and theta-gamma phase-amplitude coupling (PAC), correlating with autism severity, and repetitive behaviors (55). These findings align with current study in FXS mouse models, where KO females exhibited elevated PAC strength between gamma amplitude and theta-alpha (4–12 Hz) phases compared to WT mice, suggesting altered neural network dynamics. Another study with magnetoencephalography identified region-specific PAC abnormalities in autism, including increased midline alpha-low gamma coupling and decreased lateral coupling (56). While PAC strength correlated with poorer social communication scores across both autistic and neurotypical groups, this association was absent within the autism cohort alone, underscoring the need for further exploration of PAC’s utility as a biomarker for autism-specific clinical applications (56).

Amplitude-amplitude coupling is a method of assessing cross-frequency amplitude coupling over time (57). As mentioned earlier, studies have identified altered theta-gamma amplitude – amplitude coupling in FXS (11, 42). Similarly in our study with mice models, KO mice show inverse correlations with theta – high gamma power, compared to WT mice in both home cage and light-dark arena. Alongside, alpha – gamma correlation is nearly zero or more positive in KO mice at both ages compared to WT mice, which shows more negative or less positive correlations. Our results are consistent with what has been reported in FXS patients, where theta-gamma CFC predominates over alpha-gamma CFC in FXS compared to controls (32). These finding suggests that theta oscillations play a more prominent role than alpha oscillations in modulating gamma activity in FXS (32). Unlike alpha modulation, which typically exerts lateral inhibition on gamma activity, theta modulation may fail to suppress background gamma activity effectively. This could result in increased asynchronous neural activity and reduced signal-to-noise ratios during cognitive tasks (58). Interestingly, no sex differences were observed in theta-gamma coupling despite known differences in low-frequency oscillations and gamma activity between males and females with FXS (32).

### Theta-beta ratio

The theta-beta ratio is commonly used in neurophysiological studies to assess cognitive and attentional processes, particularly in conditions like ADHD and ASD (59, 60). Theta-beta ratio could be affected in FXS since evidence of increased theta power has been shown (11, 40). However, how theta-beta ratio is affected in FXS is unclear, although one study in FXS participants, including male and female (age range; 5 – 28 years old), did not report changes in theta-beta ratio (24). But they found it was negatively correlated with age across brain regions in both control and FXS groups (24). Alongside, they reported a positive relationship between higher theta-beta ratio (from left frontal area) and hyperactivity (24). These findings are consistent with the literature reporting an association between hyperactivity and higher theta-beta ratio in ADHD children (61).

In the current study with FXS mice, theta-beta ratio is decreased in KO mice at both ages compared to WT controls. This decrease in theta-beta ratio could be a result of elevation in beta EEG power in KO mice at both ages. Currently, there is no extensive research on the theta-beta ratio in FXS patients and rodent models. While theta-beta ratio is a well-established biomarker for ADHD, its role and significance in FXS and ASD remain unclear, highlighting the need for further research in this area.

### Peak alpha frequency

Peak alpha frequency is an EEG marker for brain maturation (62). During typical development, it shifts from theta to alpha range (62). The dynamics of neuronal oscillations in the alpha range are associated with cognition in typical development (62).

The limited available literature suggests that peak alpha frequency does not increase with chronological age in children with ASD (63). Dickinson et al. reported that peak alpha frequency decreased in ASD group compared to typically developing control children (62). Moreover, within the ASD group, peak alpha frequency correlated strongly with non-verbal cognition (62). As it reflects the integrity of neural networks, they suggested that deviations in network development may underline cognitive function in individuals with ASD (62).

A study reported that peak alpha frequency is lowered in FXS participants (male and female, age: 5 – 28 years old), ranging in the theta frequency range, rather than the alpha frequency range (33). This alteration suggests a failure in the typical maturation process where it shifts from theta to alpha frequencies during development. Such changes are consistent with findings in other neurodevelopmental disorders like ASD and ADHD, where reduced peak alpha frequency is also observed (33). Alongside, peak alpha frequency was associated with age in neurotypical control groups; however, in the FXS group, correlations were found only in central but not lateral regions of interest (33).

In our current study, peak alpha frequency was increased in KO mice compared to WT controls at juvenile age in light-dark and open field arenas, however, no change is observed in home cage recording. These results are contrasting to what is found in human FXS participants, where peak alpha frequency is lowered and shifted towards theta range (33). This discrepancy suggests that peak alpha frequency alterations observed in humans may not be fully recapitulated in the FXS mouse model. However, more future studies in rodents’ models are needed to further validate these contrasting patterns in FXS mouse models.

### Signal complexity: Multi-scale entropy

Brain signal complexity increases with age in neurotypical populations (44, 64, 65). Additionally, this increase has been shown to reflect the maturation patterns of specific sensory brain regions (43).

In current study, EEG signal complexity was found to be unchanged between WT and KO mice at both ages as indicated by complexity index at MSE (1-40). Alongside, no differences were observed at lower (1-20) and high (21-40) MSE scales as well. Our results are contrasting to what is reported in FXS patients, where EEG signal complexity was significantly lowered in FXS participants compared to healthy controls. This reduction aligned with the disruptions in brain maturation and the developmental delays associated with FXS (24).

Although findings are somewhat inconsistent, multiple studies involving individuals with ADHD and ASD have generally reported a reduction in EEG signal complexity compared to control groups (45-47). Furthermore, one specific study identified decreased complexity within the alpha frequency band in individuals with ADHD (48).

The discrepancy between human and rodent studies suggests that signal complexity alterations observed in humans may not be fully recapitulated in the mouse model.

### EEG phenotypes in FXS patients and rodents’ models

For EEG to be a reliable biomarker, it is crucial to determine its phenotypic consistency across different factors such as sexes and developmental stages in various study models.

Adding to previous studies, our results here indicated that increased gamma power in both absolute and relative EEG signal was consistent across patients and rodent models in both sexes (11, 20-23, 40, 66-68).

Regarding females, one study reported that adult FXS patients showed increase in relative theta power (11). In contrast, rat models reported a decrease in relative theta power (23, 68). Similarly, in the current study we also found a decrease in relative theta power. Thus, it seems to be opposite changes in the theta frequency range between female FXS patients and rodent models. However, it should be noted that research on female FXS patients is limited, with relatively small sample sizes (11). Alongside, a few studies have mentioned that neurotypical women exhibit greater resting state total EEG power compared to neurotypical men (69-71). Such gender-specific variations in EEG phenotypes are clinically significant, indicating potential differences in the impact of low-frequency modulation on gamma power between genders (26). Overall, there is a pressing need for further research involving female FXS patients to enhance our understanding of how gender, symptom severity, and changes in EEG interrelate.

Regarding sex difference in FXS mouse models, male KO mice showed an increase in absolute gamma EEG power, and no change in other frequency bands at both juvenile and adult ages, in both FVB and B6 background (20, 22, 67). In our current study with female mice, juvenile KO mice on the FVB background showed an increase in absolute alpha, beta and gamma EEG power. Therefore, we conclude that there is potentially sex-based EEG phenotypic difference in juvenile and adult mice with FVB background. Interestingly, one study on adult KO female mice on the B6 background reported no change in absolute or relative gamma EEG power (22). Therefore, while choosing the FXS mouse model, we should consider the background strain of mice, as well as their sex and developmental stage, as EEG phenotypes could depend on these variables.

## Author’s contribution

AA participated in EEG surgery, recording, data analysis and interpretation, and drafted and revised the manuscript. VR participated in EEG surgery, recording, data analysis and interpretation. MB participated in EEG recording. KM participated in data analysis and interpretation. NC designed the study, participated in data analysis and interpretation, and revised the manuscript. All authors read and approved of the final manuscript.

## Funding statement

This work was supported by the Alberta Children’s Hospital Research Foundation (NC), University of Calgary Faculty of Veterinary Medicine (NC), Natural Sciences and Engineering Research Council of Canada (NC), and FRAXA Research Foundation (NC). The funding sources had no role in the study design; in the collection, analysis, and interpretation of data; in the report’s writing; and in the decision to submit the article for publication.

## List of Abbreviations Used (if any)

FXS: Fragile X Syndrome
FMR1: Fragile X Messenger Ribonucleoprotein 1
FMRP: Fragile X Messenger Ribonucleoprotein 1
EEG: Electroencephalography
KO: Knockout
WT: Wildtype
PAC: Phase-amplitude coupling
AAC: Amplitude-amplitude coupling
PAF: Peak alpha frequency
TBR: Theta-beta ratio
MSE: Multi-scale entropy

## Notes

### Competing Interest Statement

The authors have declared no competing interest.

### Summary of Updates

We have now included data from adult mice (P70) in addition to our previous analyses. Furthermore, we have expanded our EEG analysis to incorporate additional parameters, including theta-beta ratio, amplitude-amplitude coupling, peak alpha frequency, and EEG signal complexity. This revised manuscript presents a comprehensive EEG study in a female fragile X syndrome mouse model at both juvenile and adult ages.

